# Connective tissue growth in a mouse model of Kosaki overgrowth syndrome is limited by STAT1

**DOI:** 10.64898/2026.04.09.717535

**Authors:** Jang H. Kim, Hae Ryong Kwon, William L. Berry, Lorin E. Olson

## Abstract

Mutations in platelet-derived growth factor receptor beta (PDGFRβ) cause Kosaki overgrowth syndrome (KOGS). Patients exhibit increased linear growth, craniosynostosis, and thin skin with increased elasticity and scarring. Of the KOGS patients identified to date, three unrelated individuals carried a P584R mutation in the juxtamembrane domain of PDGFRβ, resulting in constitutive receptor activation. Due to the limited number of patients, extensive phenotyping and exploration of the molecular basis of disease, including modifier genes, has not been completed. We generated conditional knock-in mice to express mouse PDGFRβ with a P583R mutation, corresponding to human P584R, under control of the endogenous *Pdgfrb* gene. Mutant mice were born at the expected ratio and appeared normal at birth. At 3 weeks of age, mutants began to exhibit connective tissue changes: increased body weight and bone length, craniosynostosis, ectopic bone in the tail and tendons, thin lipodystrophic skin, and high incidence of penile and rectal prolapse. To identify signaling changes caused by mutant PDGFRβ signaling, we performed western blotting and phosphoproteomics on dermal fibroblasts. This uncovered increased phosphorylation of PDGFRβ, PLCγ, Akt1, Shp2, STAT1, STAT2, STAT3, and STAT5. Analysis of 6,621 proteins and 5,386 phosphopeptides identified upregulation of interferon signaling genes linked to STAT1. In many cell types, STAT1 has tumor-suppressor functions and acts to inhibit cell cycle. We generated *Stat1*^-/-^ *Pdgfrb*^+/P583R^ mice to test the contribution of STAT1 to KOGS phenotypes. *Stat1*-deletion exacerbated overgrowth and calvaria dysmorphogensis, and caused keloid-like skin fibrosis. No phenotypes present in the original *Pdgfrb*^+/P583R^ mice were reverted to normal after *Stat1* deletion. Therefore, the P583R mouse model mirrored KOGS phenotypes and increased activation of multiple PDGFRβ signaling mediators; in this context, STAT1 activity opposes PDGFRβ-driven overgrowth and fibrosis.

## INTRODUCTION

Kosaki Overgrowth Syndrome (KOGS) is an ultra-rare syndromic disease caused by mutations in *PDGFRB*, which encodes platelet-derived growth factor receptor-β (PDGFRβ). The KOGS clinical phenotype is characterized by tall stature, scoliosis, coarse facial features, craniosynostosis, and thin skin with hypertrophic scars^1,2^. Cardiovascular phenotypes have also been identified, including progressive arterial dilation, aneurysms and heart valve defects^3-5^. KOGS was first associated with a P584R point mutation in *PDGFRB* (OMIM 616592), but it has been expanded to include several other variants^4,6,7^. These mutations lead to constitutive PDGFRβ signaling and activation of downstream signaling pathways^6,8^. Preliminary cases suggest that PDGFRβ-selective tyrosine kinase inhibitors can be used to treat KOGS^3,8,9^. However, the cellular and molecular mechanism by which mutant PDGFRβ signaling causes these phenotypes is unknown.

PDGFRβ is a receptor tyrosine kinase that plays essential roles in mesenchymal cell differentiation, proliferation, and migration^10,11^. In adult tissues, it is expressed by stromal-vascular cell types including smooth muscle cells, pericytes, fibroblasts and osteoblasts. PDGFRβ is highly conserved between mouse and humans with 95% amino acid identity in the intracellular domain. Upon PDGF ligand binding, two PDGFRβ subunits dimerize and activate their tyrosine kinase domains, leading to auto-phosphorylation. When ligand is absent, the receptor folds into an auto-inhibited conformation involving contact between the juxtamembrane and kinase domains^12^. Mutations like P584R, located in the juxtamembrane domain, disrupt auto-inhibition and lead to constitutive kinase activity^12,13^. Major downstream mediators of PDGFRβ signaling include phosphatidylinositol 3-kinase (PI3K), phospholipase Cγ (PLCγ), the tyrosine phosphatase Shp-2, Src tyrosine kinase, and adaptor proteins such as Shc and Crk^14,15^. It has been reported that Akt1, PLCγ, and Shp-2 phosphorylation are increased in KOGS^8^. Several members of the signal transducer and activator of transcription (STAT) family, namely STAT1, STAT3 and STAT5, can also be activated by PDGFRβ^16,17^. STAT1 is constitutively phosphorylated in KOGS^8^. Therefore, it is plausible that overgrowth in KOGS patients is due to a deviation from normal PDGFRβ signaling through hyperactivation of pro-growth signaling pathways.

Besides KOGS, other mutations in PDGFRβ have been linked to different human diseases^18^. Previously we characterized PDGFRβ knockin mice expressing a D849V mutation, corresponding to D850V in humans, a mutation that causes infantile myofibromatosis^19,20^. Like P584R, this mutation in the kinase domain generates constitutive PDGFRβ signaling. D849V mice developed lethal autoinflammation by 2-weeks-old due to hyperactivation of interferon signaling genes. Knockout of *Stat1* rescued autoinflammation and allowed *Pdgfrb*^*+/D849V*^*Stat1*^*-/-*^ mice to survive and develop connective tissue overgrowth^21,22^. Although many PDGFRβ mutations can lead to STAT1 activation, it is unclear whether STAT1 has the same pro-inflammatory and anti-growth properties in all contexts.

To better understand the molecular and cellular pathologies underlying KOGS, we created a mouse model to express a mutant form of mouse *Pdgfrb* corresponding to human P584R. These mice recapitulated many key phenotypes of human KOGS. To understand signaling changes due to the *Pdgfrb* mutation, we used a phosphoproteomics approach with control and mutant mouse fibroblasts. This predominantly identified differences related to interferon signaling, catabolic processes, and other gene sets linked to STAT1. Deletion of *Stat1* led to more severe overgrowth phenotypes. We conclude that in the context of PDGFRβ-related growth control, STAT1 activity moderates overgrowth by suppressing pro-growth pathways downstream of PDGFRβ.

## RESULTS

### Generation and characterization of the P583R mouse model

We generated the mouse model by replacing one *Pdgfrb* allele with a lox-STOP-lox-*Pdgfrb* cassette containing mouse *Pdgfrb* cDNA with a P583R mutation (corresponding to the human P584R). This established a line of mice where the mutation is regulated by the endogenous *Pdgfrb* gene in Cre/lox-dependent fashion (*Pdgfrb*^*LSL-P583R*^) (Figure 1A). We crossed the *Pdgfrb*^*LSL-P583R*^ allele with epiblast-expressed *Sox2-Cre*^*tg*^ to generate mice where the mutation is expressed in early embryogenesis, as occurs in most humans with KOGS. The resulting mutant mice, hereafter called P583R mice (*Sox2-Cre*^*tg*^*Pdgfrb*^*LSL-P583R/+*^), were born at the expected Mendelian frequency and were outwardly indistinguishable from wild type (WT) littermates (*Sox2-Cre*^*tg*^*Pdgfrb*^*+/+*^).

**Figure 1.**
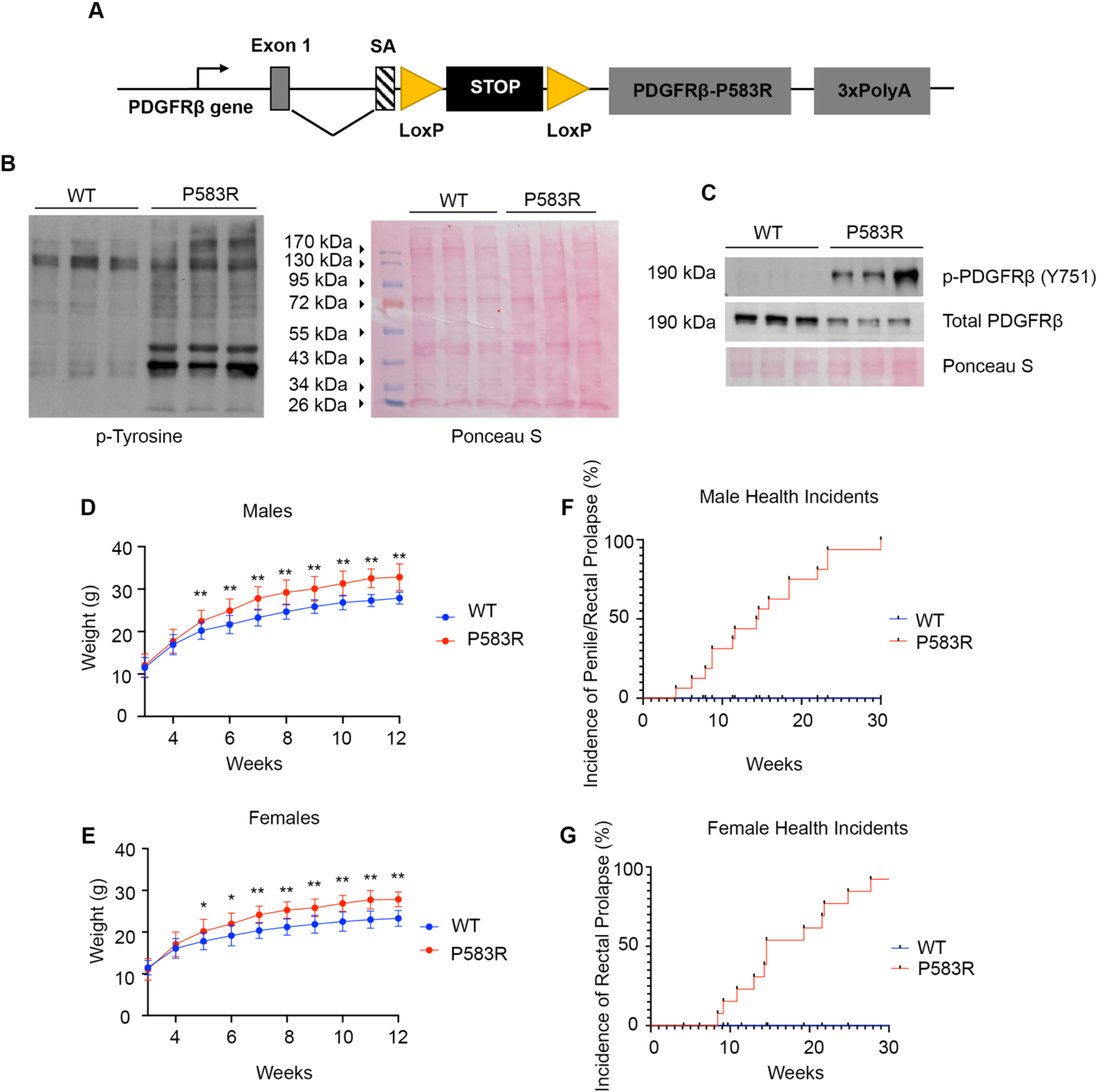
Generation and characterization of the P583R mouse model. (A) Schematic of the P583R knock-in allele for Cre-inducible expression of P583R-mutant *Pdgfrb* cDNA under control of endogenous *Pdgfrb* promoter. SA = splice acceptor. (B) Left side is western blot for phospho-tyrosine in serum-starved dermal fibroblasts. Right side = Ponceau S staining of same membrane to show protein loading in each lane. (C) Western blot for total and phosphorylated PDGFRβ in serum-starved dermal fibroblasts. (D) Growth curves of P583R males from 3-12 weeks, n=14-19 mice per genotype. (E) Growth curves of P583R females from 3-12 weeks, n=10 mice per genotype. (D-E) Error bars indicate mean +/-SD, unpaired Student’s t-test, * P < 0.05, ** P < 0.01. (F) Incidence of penile or rectal prolapse in males. (G) Incidence of rectal prolapse in females.

To verify constitutive activity of P583R, we isolated newborn dermal fibroblasts (NDFs) for western blotting. Overall tyrosine phosphorylated proteins were increased in P583R compared to WT after serum starvation (Figure 1B). As expected, P583R also displayed an increase in phosphorylated PDGFRβ, but decreased total PDGFRβ (Figure 1C).

Before 3 weeks old, P583R mice were indistinguishable from WT littermates. Older P583R mice displayed an increase in weight compared to WT, with males and females becoming significantly heavier at 5 weeks (Figure 1D-1E). Both sexes of P583R mice developed prolapses that required euthanasia. Males developed penile and rectal prolapse starting as early as 4 weeks and reaching 100% penetrance by 30 weeks (Figure 1F). Females developed rectal prolapse as early as 8 weeks and reached 100% penetrance by 30 weeks (Figure 1G). Prolapses occur in mice for many reasons and often go unexplained, but it is unusual to occur so consistently in young mice. Contributing factors may include inflammation, obstruction and straining, or defects in supporting muscle and fascia. To summarize initial characterization, the P583R mice mirrors the progressive overgrowth of KOGS, but unexpectedly they also display recurrent prolapses of unknown etiology.

### P583R mice develop skeletal overgrowth

P583R mice were not obese, but their increased weight correlated with an increase in the size of the skeleton in both sexes. WT and P583R skeletons were the same size at postnatal day 1 (Supplementary Figure 1A). P583R skeletons were larger by 12 weeks, with thicker vertebrae and barrel-shaped ribcages (Figure 2A). At birth, P583R skulls were the same size, with sutures clearly seen between mineralized calvaria (Supplementary Figure 1B). However, by 15 weeks, P583R skulls were overgrown and all cranial sutures were prematurely fused, indicating craniosynostosis (Figure 2B and Supplementary Figure 2A). Adult P583R tibias and forelimbs were longer than WT (Supplementary Figure 1C-D). Three-dimensional reconstruction of microcomputed tomography scans showed increased trabecular bone area in P583R tibias at 15 weeks (Figure 2C). P583R trabecular bone increased in total volume (TV), bone volume (BV), and trabecular number (Figure 2D-F) with an accompanying decrease in trabecular thickness, separation and mineralization (Figure 2G-I). Females were more mildly affected with only increased bone volume and trabecular number accompanied by decreased trabecular separation (Supplementary Figure 2B-H). Interestingly, cortical bone was affected in the opposite direction, with decreased total volume and bone volume in males and females (Figure 2J-N, Supplementary Figure 2I-M). Aditionally, medullary area was dramatically reduced in females. Trabecular and cortical bone growth are regulated by different mechanisms^23^. The growth plate chondrocytes generate trabecular bone and drive longitudinal bone growth, both of which were increased in P583R tibias. On the other hand, periosteal and endosteal osteoblasts generate cortical bone and circumferential bone growth, which were decreased in P583R tibias. These data suggests that the P583R mutation promotes growth plate activity but the effect on periosteal/endosteal cells is more complex.

**Figure 2.**
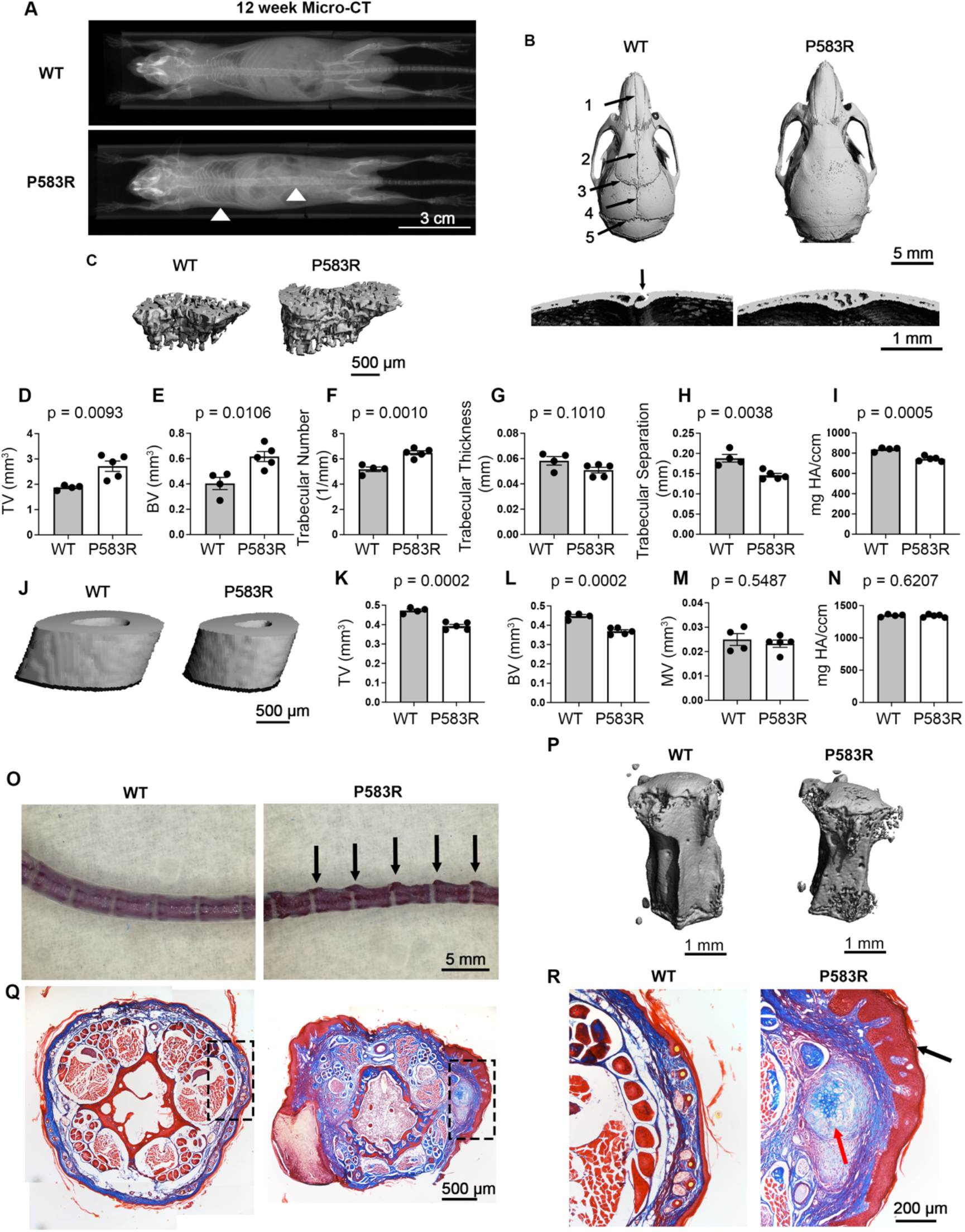
Skeletal overgrowth in P583R mice. (A) Whole body X-ray of 12-week-old WT and P583R mice. Arrowheads indicate enlarged ribcage and thickened spine. (B) μCT images of male skulls at 15 weeks showing craniosynostosis of all P583R cranial sutures. Arrows indicate sutures: 1 = internasal suture, 2 = interfrontal suture, 3 = coronal suture, 4 = sagittal suture, 5 = lambdoidal suture. Arrow in the cross section view indicates patent lambdoidal suture in WT, which is closed in P583R. (C) 3D reconstruction from μCT scans of 15-week tibias showing trabecular bone. (D) Trabecular bone total volume. (E) Bone volume of trabecular bone. (F) Trabecular number. (G) Trabecular thickness. (H) Trabecular separation. (I) Trabecular bone mineral density. (J) 3D reconstruction from μCT scans of 15-week tibias at the point of fibula attachment. (K) Cortical bone total volume. (L) Bone volume of cortical bone. (M) Cortical bone medullary volume. (N) Cortical bone mineral density. (O) Whole-mount Alizarin Red staining of tails at 26 weeks. Arrows indicate enlargement of the caudal vertebrae transverse processes. (P) 3D reconstruction of mouse tail vertebrae from μCT analysis at 15 weeks. (Q) Masson’s trichrome staining of tail lesions of P583R mice at 26 weeks. Dotted box indicates area enlarged in (R). (R) Zoom in showing ectopic cartilage nodule (red arrow) and hyperkeratotic epidermis (black arrow). Error bars indicate mean +/-SEM, unpaired Student’s t-test, n = 4-5 mice per genotype, each dot represents one mouse. Panel A shows female mice; all other panels are from male mice.

The proximal tails of P583R mice developed regularly spaced callus-like lesions by 15 weeks (Figure 2O). Tail vertebrae reconstructions showed preservation or enlargement of the vertebrae ends with narrowing of the cortical midsections, reminiscent of altered tibia dimensions (Figure 2P). Histological analysis revealed that soft tissue in the P583R tail was sclerotic, with increased collagen area, smaller muscle bundles, and almost no fat (Figure 2Q). P583R paws were also sclerotic (Supplementary Figure 1H-I). Changes in the tail and paws are interesting because some PDGFRβ mutations are temperature sensitive^24^, which might explain distinctive lesions on cooler parts of the body. Isolated P483R tail tendons were thin and dystrophic (Supplementary Figure 1E). The tail lesions themselves were hyperkeratotic and spaced over transverse processes of the vertebrae and sometimes overlying nodules of cartilage (Figure 2R). Ectopic bone formation in soft tissue, known as heterotopic ossification, is caused by genetic disorders or trauma. P583R mice exhibited heterotopic ossification in the Achilles tendon at 9 weeks (Supplementary Figure 1F-G).

### Progressive dermal atrophy and lipodystrophy in P583R mice

The skin of P583R mice was indistinguishable from WT at 2 weeks, exhibiting anagen hair follicles and a thick layer of dermal fat (Figure 3A-B). However, by 15 weeks, P583R skin became progressively thinner, with a significant reduction in upper dermis and almost no dermal fat (Figure 3C-D). Immunostaining for perilipin 1 (Plin1) confirmed a reduction in dermal adipocyte area (Figure 3E). Adipocyte gene expression was significantly reduced in P583R skin (Figure 3F). This skin atrophy phenotype resembles what we previously described in 2-weeks-old *Sox2-Cre*^*tg*^*Pdgfrb*^*+/D849V*^ mice^19^, although here it occurred with delayed onset. In the previous mice, skin atrophy was caused by PDGFRβ-driven STAT1 activity, and the phenotype could be reversed by deletion of *Stat1*^*21*^. To determine whether similar STAT1 activity could be present in P583R mice, we examined expression of STAT1-dependent interferon signaling genes (ISGs). By qPCR, ISGs were significantly overexpressed in P583R skin compared to WT (Figure 3G). Thus, the P583R mouse skin phenotype mirrors the thin skin and lipodystrophy seen in KOGS. In light of previous work with D849V mice, overexpression of ISGs suggests that STAT1 hyperactivation may cause thin skin in P583R mice.

**Figure 3.**
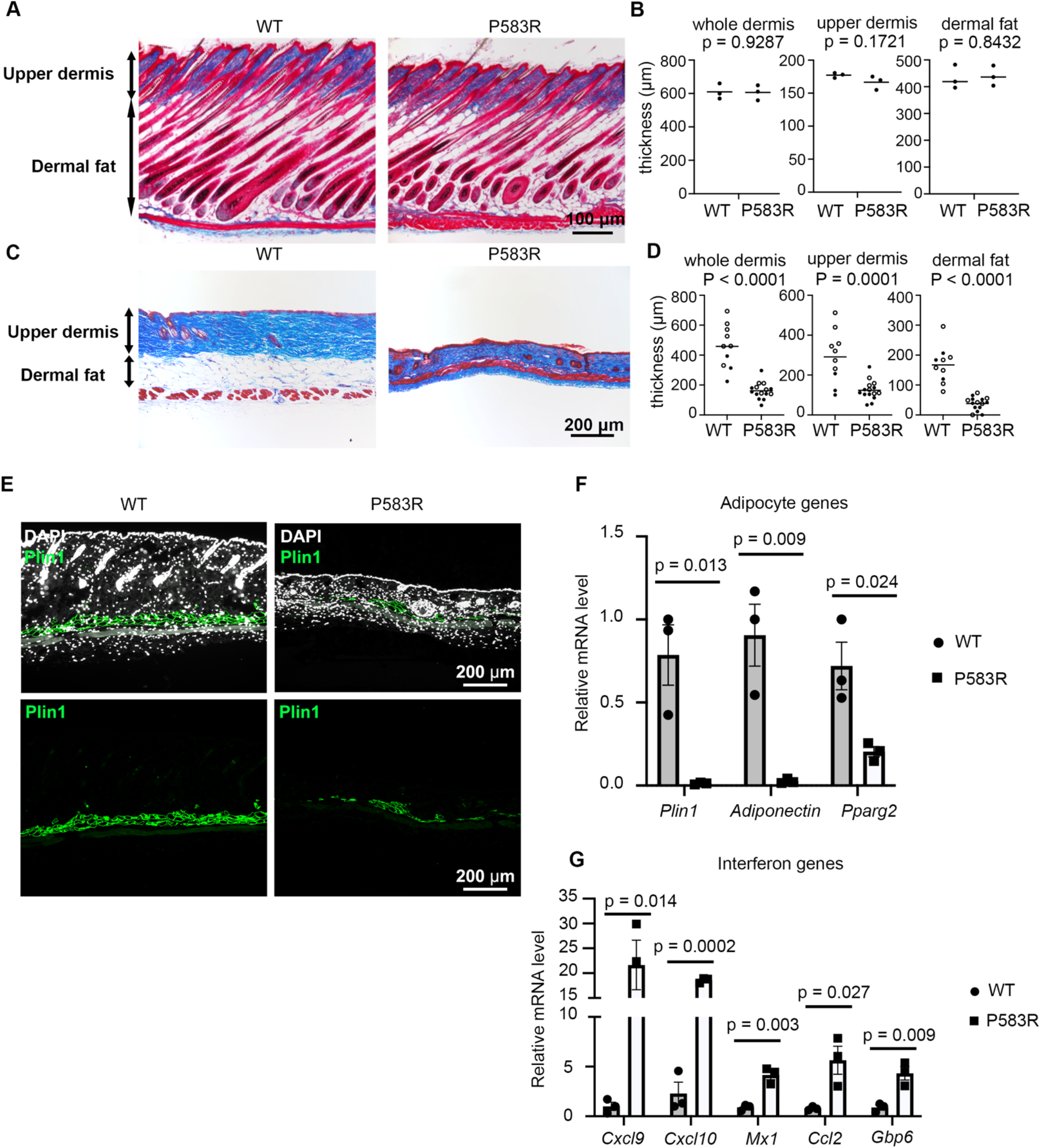
Progressive dermal atrophy and lipodystrophy in P583R mice. (A) Trichrome staining of dorsal skin at 2 weeks. (B) Quantifications of whole dermis, upper dermis, and dermal fat thickness at 2 weeks, n = 3 mice per genotype. (C) Trichrome staining of dorsal skin at 15 weeks. (D) Quantifications of whole dermis, upper dermis, and dermal fat at 15 weeks, n= 10 WT, 15 P583R. White and black circles represent males and females, respectively. (E) Immunofluorescence staining of Plin1 at 15 weeks. (F) qPCR of 15-week skin for expression of adipocyte genes. (G) qPCR of 15-week skin for expression of interferon signaling genes. Error bars indicate mean +/-SEM, unpaired Student’s t-test, n = 3 WT and 3 P583R for adipocyte genes and interferon signaling genes. Each dot represents one mouse.

### P583R constitutively activates PDGFRβ kinase cascades and activates STATs independently of JAK1/2

To examine signaling pathways, we performed western blotting on lysate from serum-starved NDFs from WT and P583R mice. AKT p-T308 phosphorylation was increased, which is a substrate for PDK1 in response to growth factor signaling. But there was little change in AKT p-S473, which is phosphorylated by mTORC2. There was no change in S6 p-S235/236, a readout of mTORC1 activity (Figure 4A). There was increased PLCγ p-Y783 and Shp2 p-Y542 in P583R cells, but no change in ERK1/2 p-T202/204 (Figure 4B). We also observed increased STAT1 p-Y701, STAT2 p-Y690, STAT3 p-Y705, and STAT5 p-Y694. For STAT1 and STAT2, even total protein was increased in P583R cells (Figure 4C), which can be explained by a positive feedback loop whereby phosphorylated STAT1 promotes its own expression and that of STAT2^25^. From these data, a pattern emerged where phosphorylation proximal to PDGFRβ (PLCγ, Shp2, AKT p-T308, STATs) were increased in P583R cells, but distal phosphorylation (ERK, AKT p-S473, S6) was not increased, potentially due to negative feedback (Figure 4D).

**Figure 4.**
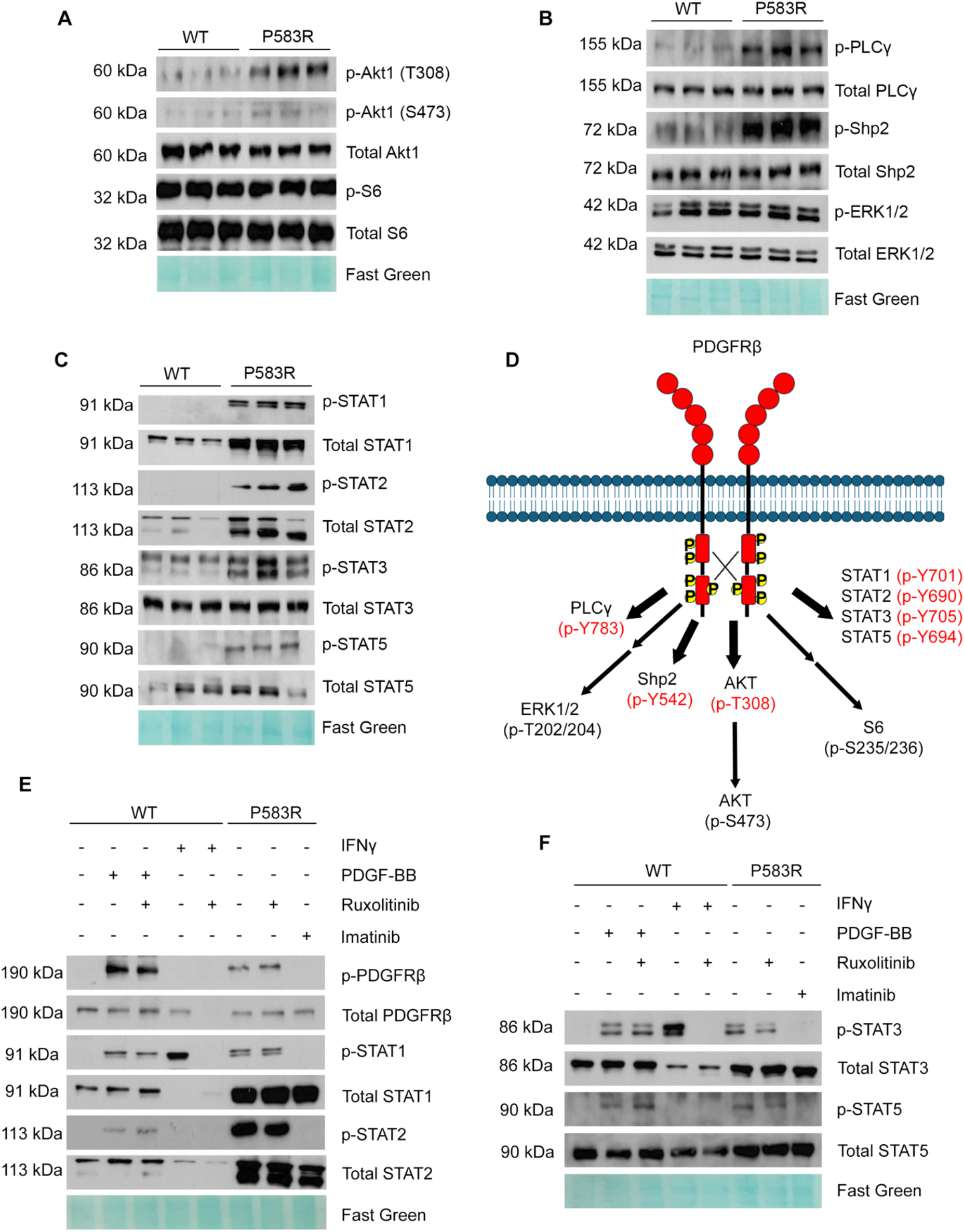
P583R activates multiple kinase cascades and STATs. (A-C) Western blotting with cell lysate from serum-starved dermal fibroblasts. (D) Cartoon schematic of distal and proximal phosphorylation events relative to PDGFRβ. Thick single arrows indicate phosphorylation by PDGFRβ or kinases close to PDGFRβ. Thin arrows indicate phosphorylation by a kinase cascade. (E-F) Western blotting on cell lysate from serum-starved fibroblasts, with pre-treatment with ruxolitinib (JAK1/2 inhibitor) or imatinib (PDGFRβ inhibitor), and stimulated with IFNγ or PDGF-BB for 15 minutes. Ruxolitinib does not block PDGF or P583R-induced STAT phosphorylation. All western blotting experiments were performed at minimum three separate times with separate biological replicates.

It is unclear whether PDGFRβ and other growth factor receptors can phosphorylate STATs directly. Cytokine receptors lack intrinsic kinase activity and recruit Janus Activated Kinases (JAKs) to tyrosine phosphorylate STAT proteins^26^. To address whether wild type PDGFRβ or the P583R variant requires JAK1/2 for STAT phosphorylation, we treated WT and P583R NDFs with ruxolitinib, a competitive inhibitor of the ATP-binding site of JAK1/2^27^. In WT treated with PDGF-BB, 1 μm ruxolitinib failed to block phosphorylation of STAT1, STAT2, STAT3 or STAT5. However, ruxolitinib blocked IFNγ-induced phosphorylation of STAT1 and STAT3; IFNγ does not activate STAT2 or STAT5 (Figure 4E-F). In P583R cells, ruxolitinib did not block any STAT phosphorylation. On the other hand, inhibition of PDGFRβ kinase activity with 1 μm imatinib was sufficient to block phosphorylation of all STATs in P583R cells (Figure 4E-F). These results show that WT and P583R forms of PDGFRβ activate STATs in a JAK-independent fashion. As a consequence of this mechanism, PDGFRβ-induced STAT activation might be resistant to negative feedback mechanisms that depend on inhibition of JAK activity, for instance via suppressors of cytokines signaling (SOCS)^28^.

### Phosphoproteomics identifies upregulated tyrosine phosphorylation and interferon signaling

To globally survey changes in protein expression and phosphorylation, we performed phosphoproteomics on serum-starved WT and P583R NDFs at passage one. We used tandem mass tag labeling to individually label 3 WT and 4 P583R biological replicates. After trypsin/Lys-C digestion, phosphopeptides were enriched with TiO_2_ and Fe-NTA, then total and phosphoproteins were identified by liquid chomatography-mass spectrometry (LC/MS). Across all samples, a total of 6,621 unique proteins and 5,386 unique phosphopeptides were detected. 98.1% of the phosphorylation events were on serine/threonine while 1.9% were on tyrosine, which reflects the relative rarity of tyrosine phosphorylation. Principal component analysis showed clear separation between WT and P583R samples for both phosphopeptides and total proteins (Figure 5A-B). Normalization of the log_2_ intensity showed that average expression and total expression of phosphopeptides and total proteins were comparable between WT and P583R (Supplementary Figure 3A-D).

**Figure 5.**
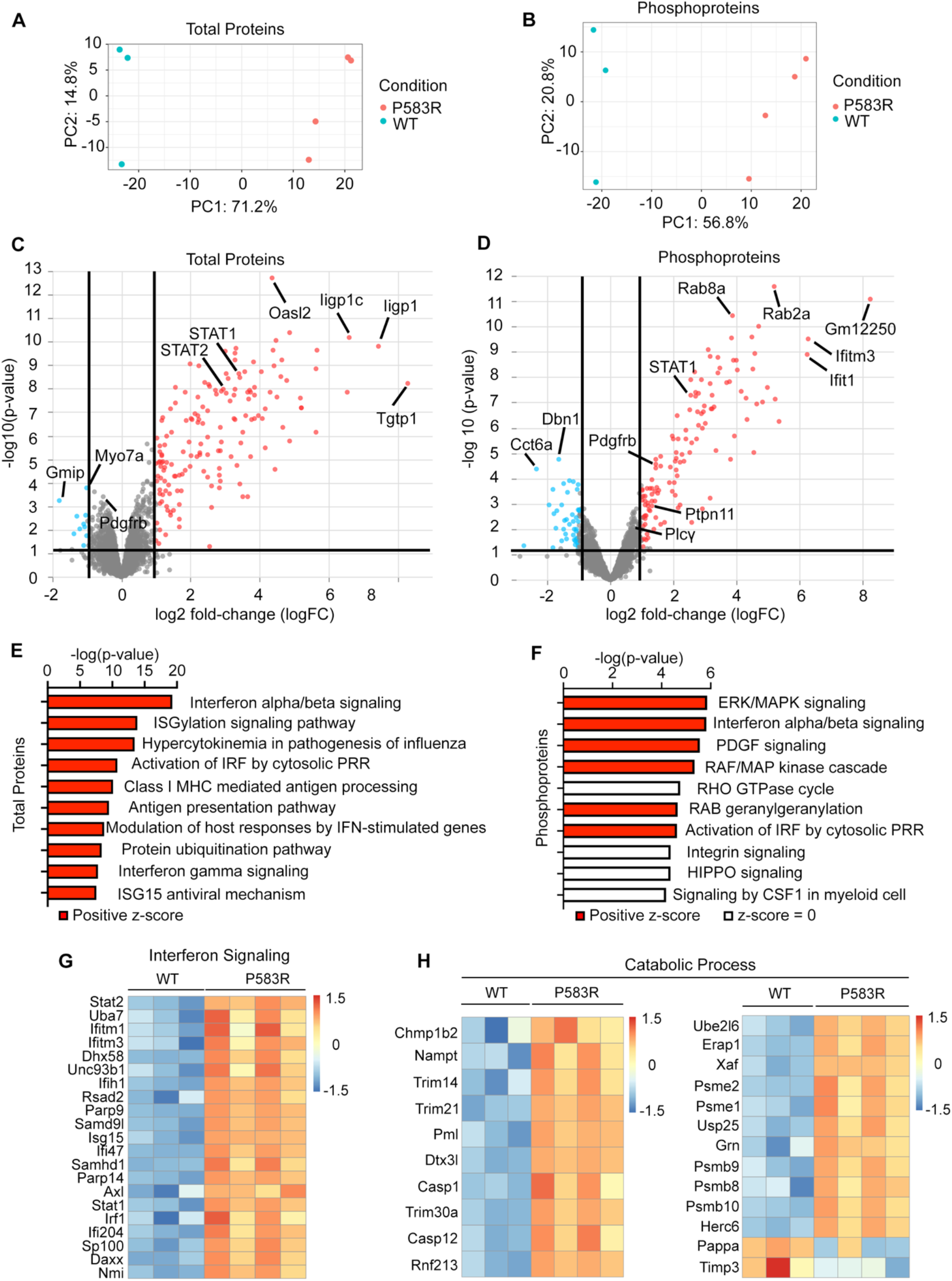
Phosphoproteomics identifies upregulated interferon signaling. (A-B) Principal component analysis based on all detected proteins or phosphopeptides. (C-D) Volcano plot of total proteins or phosphorylated peptides. (E-F) Ingenuity Pathway Analysis of significantly differentially expressed proteins or phosphopeptides showing enrichment of pathways. Positive z-score suggests activation of a pathway. Z-score of zero suggests changes without clear activation or deactivation of a pathway. (G) Differentially expressed proteins comprising “Interferon Signaling” category. (H) Differentially expressed proteins comprising “Catabolic Process” category. Significance cutoff for proteins and phosphopeptides is log_2_ intensity > 1 and p-value < 0.05.

Volcano plots depicted 167 differentially expressed proteins (DEPs) in P583R with most (n = 124) showing increased expression (Figure 5C). In agreement with the western blots, STAT1 and STAT2 were significantly increased in P583R. PDGFRβ was slightly decreased but did not reach statistical significance. There was differential phosphorylation of 170 peptides with most (n = 160) being increased (Figure 5D). Among these, STAT1 p-S727, Shp2 p-Y542, PDGFRβ p-Y751, and PDGFRβ p-Y856 were significantly increased while PLCγ p-Y783 was slightly increased but not significantly. Other phosphoproteins identified by western blotting (Akt, S6, Erk) were not detected in the phosphoproteomics dataset.

As mentioned earlier, peptides with tyrosine phosphorylation, which could be direct targets of PDGFRβ, were just 1.9% of the 5,386 phosphopeptides detected. However, 28.2% (n = 48) of the 170 differentially phosphorylated peptides were tyrosine phosphorylated. We used the Kinase Library^29,30^ to identify 14 phosphotyrosines with recognition sequences predicted as substrates for PDGFRβ kinase. This short list included known PDGFRβ substrates: PDGFRβ itself, Vimentin, Ptpn11 (encoding Shp2), Shc1, PLCγ, and Ephb2. There were also proteins that could reasonably interact with PDGFRβ at the plasma membrane: AnnexinA2, Ephb3, Dlg1, Hgs, and Pard3. There were three proteins with PDGFRβ-like recognition sequences that are not obviously related to PDGFRβ or membrane localization: G6pdx, Zdhhc3, Pin4 (Supplementary Figure 3E).

To identify biological process associated with the DEPs or differentially phosphorylated proteins (DPPs), we used Ingenuity Pathway Analysis. Based on DEPs, the top ten upregulated pathways comprised processes related to interferon signaling (Figure 5E), which were all linked by STAT1 (Supplementary Figure 4A). Based on phosphorylation differences, the top upregulated pathways included ERK/MAPK, interferon, and PDGF signaling cascades (Figure 5F). Following gene set enrichment analaysis, heatmaps in Figure 5G-H show DEPs related to interferon signaling and catabolic processes involving ubiquitination and proteasome activity. There was also upregulation of DEPs involved in ribonucleotide binding, GTPase activity, and antigen presentation (Supplementary Figure 4B-D). Most of these DEPs are associated with STAT1-regulated processes and are considered ISGs, for example Psmb8, Psmb9, and Psmb10 (Figure 5H) are proteasome subunits of the immunoproteasome that processes antigens for MHC class I presentation. Together, these results demonstrate that the P583R variant activates PDGF-regulated kinase pathways alongside potent activation of STAT1 signaling, with STAT1 target ISGs dominating the DEPs identified by proteomics.

### Deletion of *Stat1* exacerbates skeletal overgrowth in P583R mice

In previous studies using the D849V variant, mutants died at two-weeks-old with autoinflammation. Deletion of *Stat1* was sufficient to eliminate ISG expression and rescue postnatal lethality, but subsequently the *Pdgfrb*^*+/D849V*^ *Stat1*^*-/-*^ mice developed overgrowth^21^. We concluded that, in the context of D849V, STAT1 has a dual role of driving autoinflammation and suppresing a latent overgrowth phenotype. The current P583R mice have a milder phenotype than D849V in terms of overall survival, but they similarly hyperactivate STAT1 and overexpress ISGs, leading us to hypothesize that STAT1 could play a similar dual role here.

We generated *Sox2-Cre*^*tg*^*Pdgfrb*^*LSL-P583R/+*^*Stat1*^*f/f*^ (PS1^-/-^) mice and *Sox2-Cre*^*tg*^*Stat1*^*f/f*^ (S1^-/-^) littermate controls. S1^-/-^ mice resembled wild type mice with a slight decrease in body weight. Deletion of *Stat1* did not rescue phenotypes seen in P583R mice. On the contrary, PS1^-/-^ males and females were significantly heavier than P583R mice (Figure 6A-6B). Prolapses occurred at similar frequency in PS1^-/-^ and P583R mice. Whole body X-ray from 15-week PS1^-/-^ mice showed an obvious increase in body length compared to S1^-/-^ littermates (Figure 6C and Supplementary Figure 5A) or P583R mice (Figure 2A). Besides thicker spines and barrel-shaped ribcages, PS1^-/-^ mice also exhibited scoliosis (Figure 6C and Supplementary Figure 5A). Craniosynostosis still occurred and skull overgrowth was exacerbated in PS1^-/-^ mice at 15 weeks old (Figure 6D and Supplementary Figure 5B). PS1^-/-^ skulls displayed enlarged supraorbital ridges, expanded and misshapen zygomatic arches, and overgrowth of the mandible and maxilla, resulting in prognathism and malocclusion. PS1^-/-^ calvaria also exhibited expanded bone marrow space (Figure 6D), similar to *Pdgfrb*^*+/D849V*^ *Stat1*^*-/-*^ mice described previously^21^.

**Figure 6.**
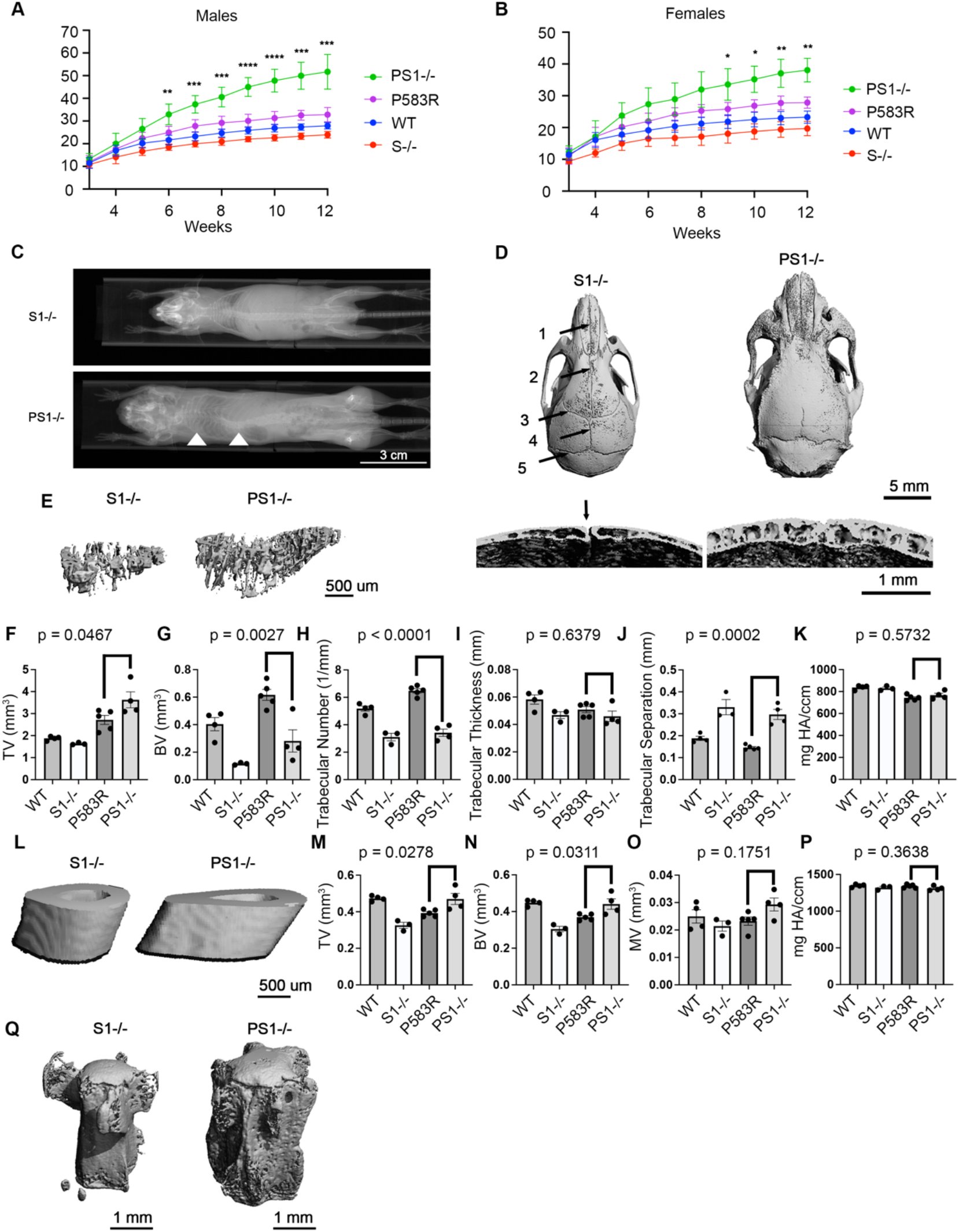
Deletion of *Stat1* exacerbates skeletal overgrowth in P583R mice. (A-B) Growth curves for wild type, P583R, PS1^-/-^ (*Sox2-Cre*^*tg*^*Pdgfrb*^*LSL-P583R/+*^*Stat1*^*f/f*^), and S1^-/-^ (*Sox2-Cre*^*tg*^*Stat1*^*f/f*^) mice. Data for WT and P583R are the same as Figure 1D-E, n = 8 male PS1^-/-^, 6 male S1^-/-^, 6 female PS1^-/-^, 3 female S1^-/-^. Error bars indicate mean +/-SD, two-way ANOVA with multiple comparisons and Tukey’s correction, * P < 0.05, ** P < 0.01, *** P < 0.001, **** P < 0.0001. (C) Whole body X-ray of 15-week-old male mice. Arrowheads indicate enlarged ribcage and scoliosis of thickened spine. (D) μCT of male skulls at 15 weeks showing overgrowth and craniosynostosis in PS^-/-^. Arrows indicate sutures. 1 = internasal suture, 2 = interfrontal suture, 3 = coronal suture, 4 = sagittal suture, 5 = lambdoidal suture. Arrow in the cross-section view indicates patent lambdoidal suture in S1^-/-^, which is closed in PS1^-/-^. (E) 3D reconstruction from μCT scans of 15-week tibias showing trabecular bone. (F) Trabecular bone total volume. (G) Bone volume of trabecular bone. (H) Trabecular number. (I) Trabecular thickness. (J) Trabecular separation. (K) Trabecular mineralization (L) 3D reconstruction from μCT scans of 15-week tibias at the point of fibula attachment. (M) Cortical bone total volume. (N) Bone volume of cortical bone. (O) Cortical bone medullary volume. (P) Mineralization of cortical bone. (Q) 3D reconstruction of mouse tail vertebrae from μCT analysis at 15 weeks. (E-P) All quantifications from male mice at 15 weeks. Data for WT and P583R are the same as Figure 2D-I, K-N. Error bars indicate mean +/-SEM, one-way ANOVA with multiple comparisons and Tukey’s correction, n = 3-5 mice per genotype, each dot represents one mouse.

Three-dimensional reconstruction of microcomputed tomography scans showed increased trabecular bone area in PS1^-/-^ tibias at 15 weeks (Figure 6E, Supplementary Figure 5C). Accordingly, trabecular total volume and trabecular separation were increased, but bone volume and trabecular number were decreased, with no change in mineralization (Figure 6F-K, Supplementary Figure 5D-I). Cortical bone was also enlarged, with increased total volume and bone volume accompanied by no change in mineralization (Figure 6L-P, Supplementary Figure 5J-N). Female medullary area was normalized in PS1^-/-^ compared to P583R (Supplementary Figure 5M). Unlike P583R mice, tail lesions were not readily apparent in PS1^-/-^ mice. However, 3D reconstructions showed that the transverse processes and cortical midsections of the PS1^-/-^ tail vertebrae were expanded (Figure 6Q). These results show that male and female skeleton overgrowth is exacerbated by the absence of STAT1.

### Keloid-like dermal fibrosis in P583R mice with *Stat1* deletion

Unlike P583R mice with lipodystrophy, skin thickness was significantly increased in PS1^-/-^ mice at 15 weeks. The phenotype involved increased fibroblasts and dense, keloid-like collagen deposition extending from the epidermis to the panniculus carnosus (Figure 7A-B). This resulted in obliteration of the dermal adipose layer, although scattered Plin1^+^ cells were still present (Figure 7C). In both sexes of PS1^-/-^ mice compared to P583R, thickness of the whole skin was increased due to expansion of the upper dermis at the expense of dermal fat (Figure 7D). Similar to P583R mice, PS1^-/-^ mice displayed significantly lower expression of adipocyte markers (Figure 7E). Consistent with being STAT1-dependent ISGs, *Cxcl9, Cxcl10, Mx1, Ccl2, Gbp6* expression was normalized compared to P583R (Figure 7E-F). Therefore, while P583R mice already exhibit overgrowth, *Stat1* deletion exacerbates the overgrowth of bone and skin, demonstrating that STAT1 plays a dual role driving autoinflammation and suppressing overgrowth in the context of P583R.

**Figure 7.**
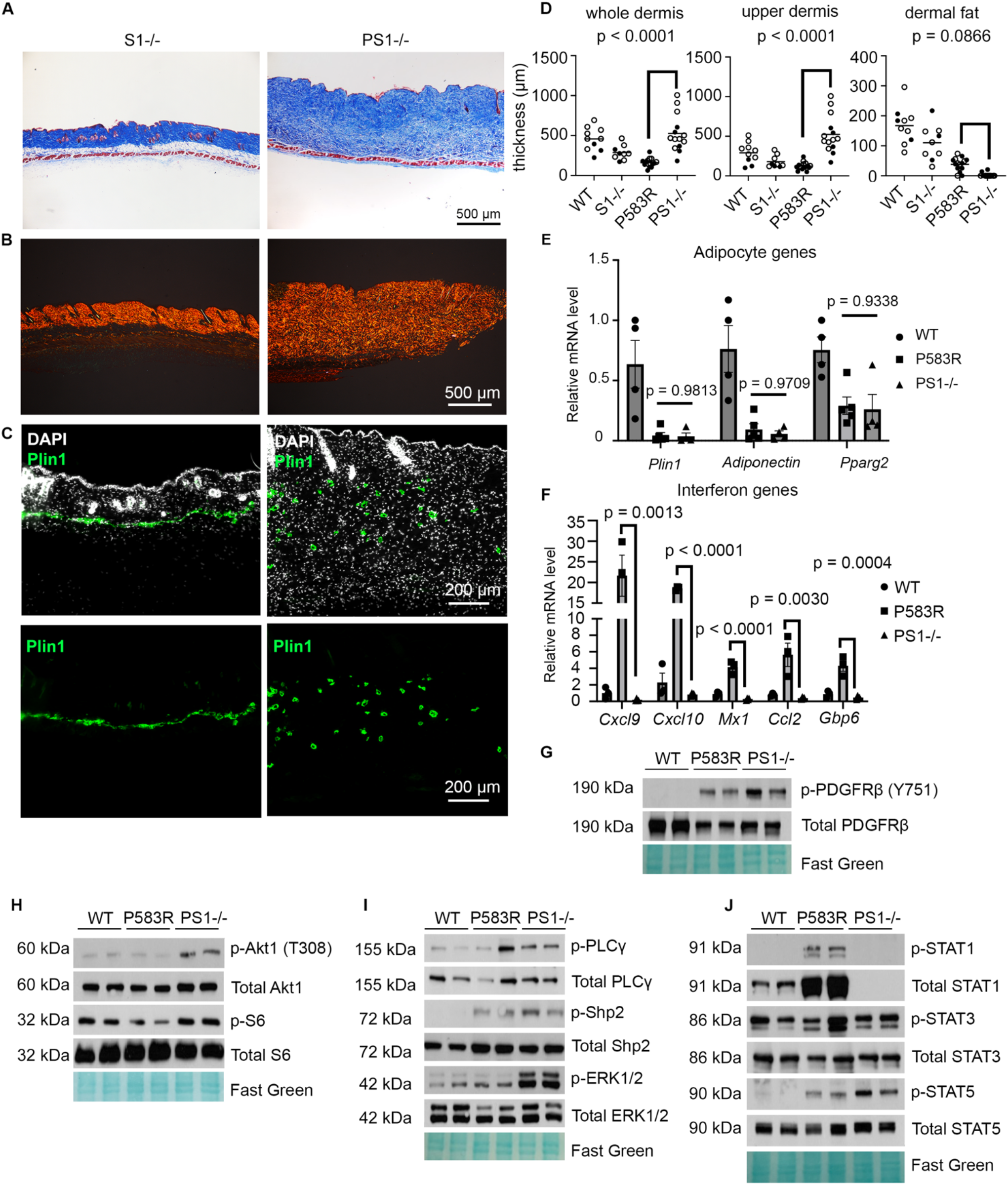
Keloid-like dermal fibrosis in P583R mice with *Stat1* deletion. (A) Trichrome staining of male skin at 15 weeks. (B) Picrosirius Red staining of dorsal skins at 15-weeks. (C) Immunofluorescence staining for Plin1 at 15 weeks. (D) Quantifications of whole dermis, upper dermis, and dermal fat at 15 weeks, n = 6 S1^-/-^ males, 3 S1^-/-^ females, 9 PS1^-/-^males, 5 PS1^-/-^ females. White and black circles represent males and females, respectively. Data for WT and P583R are the same as Figure 3D. (E) qPCR of 15-week skin for expression of adipocyte genes. (F) qPCR of 15-week skin for expression of interferon signaling genes. (D-F) Data for WT and P583R are the same as Figure 3F-G. Error bars indicate mean +/-SEM, one-way ANOVA with multiple comparisons and Tukey’s correction. n = 3-5 mice per genotype, each dot represents one mouse. (G-J) Western blotting with cell lysate from serum-starved dermal fibroblasts.

### Deletion of *Stat1* increases phosphorylated levels of ERK1/2

The dramatic worsening of skeletal overgrowth and dermal fibrosis suggested that removing STAT1 might change PDGFRβ signaling or its downstream signaling pathways. We performed western blotting on lysates of serum-starved NDFs from WT, P583R, and PS1^-/-^ mice. Total and p-Y751 forms of PDGFRβ were slightly increased in PS1^-/-^ compared to P583R, suggesting that removal of STAT1 has some effect on PDGFRβ activity (Figure 7G). Phosphorylated PLCg, Shp2 and STAT3 were not increased in PS1^-/-^ compared to P583R. There was, however, a slight increase in phosphorylated Akt1, S6, and STAT5 phosphorylation, and a prominent increase in ERK phosphorylation (Figure 7H-J). These results show that deletion of *Stat1* relieves an inhibitory signal that limits PDGFRβ-directed pro-growth pathways.

## DISCUSSION

Activating mutations in PDGFRβ have been linked to several human diseases including Kosaki overgrowth syndrome, Penttinen syndrome, and infantile myofibromatosis^18^. Distinct disease features led to their classification as different diseases, with Penttinen syndrome characterized by progeroid features, and myofibromatosis by benign fibrous tumors. However, similar tumors can occur with KOGS, and some Penttinen patients display increased growth like KOGS. It has been suggested that KOGS and Penttinen syndrome are variants of a single disease^3^. One of the questions we set out to address was whether a variant of PDGFRβ that causes KOGS would generate KOGS-like phenotypes when introduced into mice. We found that P583R mice generated with *Sox2-Cre*^*Tg*^ recapitulate major features of KOGS including progressive weight gain, skeletal overgrowth, and craniosynostosis, with somewhat later development of lipodystrophy and thinning of the skin. We have not seen tumors in P583R mice. Human patients with KOGS have dilation and tortuosity of major arteries, often leading to aneurysms and strokes at a young age^3-5^. Up until 30 weeks of age, we have not observed P583R mice to experience sudden death due to cardiovascular events. However, this question merits further investigation using Cre-drivers that specifically target vascular cell types. The skeletal overgrowth and dermal changes in this new model provide an entry point for exploration of cellular and molecular mechanisms of KOGS.

P583R causes constitutive PDGFRβ signaling, resulting in hyperactivation of kinase pathways and hyperphosphorylation of multiple STATs. We confirmed many of the signaling changes previously reported for human primary cells or cell lines expressing P584R. These include increased phosphorylation of PLCγ, Shp2, Akt, STAT1, STAT3 and STAT5. Homodimers of tyrosine phosphorylated STAT1 are the principal mediators of type II interferon signaling. We also observed increased phosphorylation of STAT2, which combines with STAT1 and IRF9 to mediate type I interferon signaling. These changes in STAT1 and STAT2 explain the strong interferon response.

In the classical JAK-STAT signaling pathway, activated cytokine receptors recruit Janus activated kinases (JAKs) to phosphorylate STAT proteins. Activated STATs translocate to the nucleus and induce STAT-specific target genes, including suppressors of cytokines signaling^26^ that provide negative feedback on JAK. Using ruxolitinib, we discovered that PDGFRβ-mediated STAT activation does not require JAK activity. This was true for wild type PDGFRβ as well as the P583R variant. This agrees with a recent finding that the V665A variant, which causes Penttinen syndrome, does not require JAK1/2 for STAT1 phosphorylation^31^. We speculate that because of this mechanism, PDGFRβ-induced STAT1 activation might be resistant to negative feedback mediated by SOCS proteins. Lacking a major negative feedback mechanism, this type of STAT1 activation could be highly damaging.

Using LC/MS phosphoproteomics, we surveyed the proteome and phosphoproteome of wild type and P583R fibroblasts to identify signaling changes in a more unbiased manner. Dermal fibroblasts were used because they are easy to isolate in large numbers from newborn skin and P583R mice develop a skin phenotype. We suspect that signaling pathway changes would be similar in skeletal cells. The P583R proteome exhibited 167 DEPs with significant enrichment of protein sets related to interferon signaling, which generally acts as an anti-growth signal in mesenchymal cells^32^. The P583R phosphoproteome included 170 DPPs, with enrichment of tyrosine phosphorylation and of signaling pathways related to PDGF, MAPK/ERK and interferon signaling. In particular, Shp2 protein (encoded by *Ptpn11*) exhibited increased tyrosine phosphorylation by western blotting and proteomics. Shp2 is interesting because it serves as a hub connecting multiple signaling pathways downstream of PDGFRβ, including MAPK/ERK activation^33,34^. Shp2 can also negatively regulate STAT1 and interferon signaling^35,36^. The combination of anti-growth and pro-growth signaling raised the possibility that STAT1 could suppress connective tissue overgrowth. This was confirmed when *Stat1* deletion unleashed further overgrowth in the PS1^-/-^ skin and skeleton. Intriguingly, STAT1 deletion slightly increased phosphorylation of PDGFRβ, Akt1, S6 and STAT5, and greatly increased phosphorylation of ERK. Further studies are required to solve exactly how removing STAT1 rewires mutant PDGFRβ kinase pathways to unleash connective tissue overgrowth.

The P583R mice bear some resemblance to the D849V mice described in previous studies^19,21,22^. A major difference is that D849V mutants died by 2 weeks old with prominent lipodystrophy, thin skin, osteopenia, and ISG overexpression. *Stat1* deletion from D849V mice rescued early lethality and, in a similar fashion to P583R mutants, unleashed overgrowth of the skeleton and skin. Since STAT1 is an important modifier gene in both models, we speculate that at least some differences between P583R and D849V phenotypes could be due to STAT1 phosphorylation being lower in P583R mutants compared to D849V^21^.

The first two KOGS patients were heterozygous for P584R variants^1^. Subsequently, KOGS has presented a range of craniofacial changes involving craniosynostosis, widening and enlargement of the nasal bridge and supraorbital ridges, and changes in dentition. Craniofacial phenotypes in P583R mice were consistent with KOGS and were exacerbated by *Stat1* deletion. The human skin phenotype has not been investigated in detail, but clinical descriptors include thin, fragile, hyperelastic and lipodystrophic skin. P583R mice exhibited a striking loss of dermal fat and skin thinning, consistent with KOGS. Deletion of *Stat1* reversed skin thinning (but not lipodystrophy) and caused fibrotic expansion of the dermis with thick bundles of collagen. Notably, a non-inducible mouse model of KOGS was generated by gene editing a W565R point mutation into the *Pdgfrb* gene, which resulted in Runx2-dependent craniosynostosis^37^. Skin phenotypes have not been described in the W565R mouse.

As P583R mice reached early adulthood, they became prone to rectal and penile prolapses requiring early euthanasia. The high incidence of prolapses was unexpected. Prolapses are unusual in young mice but become more common with age as connective tissues weaken. Other factors that contribute to prolapses include infection, inflammation, obesity, obstruction and straining, or weakness in supporting muscle and fascia. Deletion of *Stat1* did not rescue prolapses, suggesting that interferon-related inflammatory mechanisms are not the cause. We speculate that fascia development defects or accelerated weakening of connective tissue contribute to prolapses in these mice.

In summary, we created a conditional knock-in model of KOGS by introducing a P583R mutation into mouse *Pdgfrb*. These mice developed skeleton and skin phenotypes that mimic the human disease. Ectopic bone and prolapse phenotypes are novel. We identified hyperactivation of pathways related to overgrowth (ERK, Akt, STAT5) and interferon signaling (STAT1, STAT2). Abrogation of the STAT1-dependent inflammatory pathway exacerbated overgrowth, which demonstrates that STAT1 is an important gene modifier for PDGFRβ-related disease involving connective tissue overgrowth.

## METHODS

### Mice

All animal experiments were performed according to procedures approved by the Institutional Animal Care and Use Committee of the Oklahoma Medical Research Foundation. Mice were maintained on a 12 hr light/dark cycle and housed in groups of two to five with unlimited access to water and food. All strains were maintained on a mixed C57BL6/129 genetic background at room temperature. Mouse strains *Sox2-Cre* (JAX 008454)^38^ and *Stat1*^*flox*^ (JAX 012901)^39^ are available from Jackson Laboratory. To generate *Pdgfrb*^*LSL-P583R*^ mice, a *Pdgfrb* cDNA containing the P583R mutation was knocked into the *Pdgfrb* locus in embryonic stem cells using a previously described vector^19^. To monitor incidences of prolapse, veterinary health reports were noted for first indication of prolapses. As prolapses progressed, P583R mice were euthanized following the recommendations of the American Veterinary Medical Association (AVMA) guidelines. Mouse weights were recorded weekly. Both sexes were analyzed and littermates were used for comparison whenever possible.

### Histology

Skin was harvested after hair removal with cream (Nair). Skin or tail tissue was fixed with 10% neutral buffered formalin overnight at 4°C. Tissues were processed, embedded in paraffin, sectioned at 8 μm, and used for Masson’s trichrome or picrosirius red staining. ImageJ was used to measure thickness of the whole dermis (between epidermis and panniculus carnosus) and the upper dermis (between epidermis and fat, or between epidermis and panniculus carnosus when the fat layer was missing).

### Immunostaining

Tissues were harvested as described above. After fixation, skins were submerged in 20% sucrose solution for 24 - 48 hours, cryoembedded, and sectioned at 20 μm thickness. After sectioning, slides were rinsed in PBS for 5 minutes to rehydrate tissue and remove excess freezing medium. Tissues were blocked with 5% serum + 0.1% Triton-X in PBS solution for 30 minutes at room temperature in a hydrated chamber. Tissues were then incubated overnight at 4°C with primary antibody solution containing 0.05% triton and 5% serum. After overnight incubation, slides were washed in PBS for 10 minutes, repeated three times. Tissues were incubated for one hour in a humidified chamber using secondary antibody solution containing 0.05% triton and 5% serum. Tissues were stained with PBS + DAPI for 5 minutes. Tissues were washed 4 times in PBS for 5 minutes per wash and coverslipped using Fluoro Gel with DABCO (Electron Microscopy Sciences). Images were captured with a Nikon Eclipse 80i microscope connected to a digital camera.

### RNA isolation for qPCR

Skin was removed, flash frozen in liquid nitrogen, and pulverized to a fine powder. The powder was resuspended in TRIzol (Thermo Fisher) to solubilize the RNA and purify it following the manufacturer’s protocol. RNase-free water was added to resuspend the RNA pellet. cDNA synthesis was performed using PrimeScript RTase (Takara), and quantitative PCR was performed using SYBR green master mix (Bio-rad), iCycler (Bio-rad) and primer sets shown in Table S2. Expression of mRNA levels were normalized relative to *Gapdh* levels.

### Whole-mount skeletal preparation

Newborn mice between perinatal day 0 and day 2 were euthanized according to humane guidelines. Mice were then immersed in a 65°C water bath for 1 minute to remove skin, and internal organs were removed. Carcasses were then fixed in 100% EtOH for 72 h, stained with Alcian Blue for 12 h (15 mg Alcian Blue, 80 mL 95% EtOH, 20 mL Glacial Acetic acid), washed in 100% EtOH overnight, cleared in 1% KOH overnight, stained with Alizarin Red for 8 h (50 mg Alizarin Red/Liter of 1% KOH), cleared in 1% KOH overnight, and washed with 50:50 glycerol:1% KOH overnight, 80:20 glycerol: 1% KOH overnight, and stored in 100% glycerol.

Adult tails were harvested and fixed overnight in 10% neutral buffered formalin. Following fixation, tissues were decolored using the rapid bone staining fixation (RAP-fix) solution (5% formalin, 5% triton, and 2% KOH) at 42° C with shaking for 48 hours until skin became white in appearance. Tails were submersed in RAP enhancement solution (20% ethylene glycol, 5% triton, and 2% KOH) at 42° C with shaking for 24 hours for clearing^40^. Skeletal staining with Alizarin Red was performed as described above, washed with 50:50 glycerol:1% KOH overnight, 80:20 glycerol: 1% KOH overnight, and stored in 100% glycerol.

### MicroCT

Euthanized mice were placed in the VivaCT 40 microCT (Scanco Medical) and scout views were taken to visualize the mineralized skeleton depicted by whole body X-ray. Bones were freshly harvested from euthanized mice and fixed in 10% neutral buffered formalin with shaking at room temperature. Bones were scanned overnight, and 3D rendered images were generated from scans. Bones were analyzed to measure bone volume (mm^3^), total volume (mm^3^), bone volume/total volume, mean/density (mg HA/ccm), trabecular numbers (1/mm), trabecular thickness (mm), and trabecular separation (mm) with a VivaCT 40 microCT using the parameters of X-ray tube voltage of 70 kVP and X-ray current of 114 μA.

### Newborn dermal fibroblast isolation and western blotting

Newborn mice between perinatal day 0 and day 2 were euthanized according to humane guidelines. Skin was removed and floated flat on 0.25% Trypsin/EDTA overnight at 4°C, dermis side down. Then skins were transferred to a dry petri dish, epidermis side down, and dermis was lifted away. Dermis was then minced with scissors and digested with Type II Collagenase (500 U/mL) dissolved in DMEM. The solution was then incubated at 37°C for one hour with agitation every 15 minutes. Cells were filtered with a 100 μM filter and rinsed with DMEM/F12 twice to ensure removal of residual collagenase. After plating and growth until confluency in DMEM/F12 with 10% fetal bovine serum, 1% L-glutamine, and 0.5% Pen/Strep, cells were serum starved in DMEM/F12 + 0.1% FBS overnight. Western blotting was performed on fibroblasts between passage 2-5. Cells were lysed in lysis buffer (50 mM Tris at pH 7.4, 1% NP-40, 0.25% sodium deoxycholate, 150 mM NaCl, 0.1% sodium dodecyl sulfate) with the addition of 1 mM NaF, 1 mM Na_3_VO_4_, 1 mM PMSF, 1mM EDTA, and 1x protease inhibitor cocktail (Complete, Roche). Fibroblast lysate was sonicated for 10 minutes, and protein concentration was determined by Pierce BCA assay. 20 μg of protein was separated by SDS-PAGE and transferred onto a nitrocellulose membrane. After transfer, staining with 0.5% Ponceau S or 0.1% Fast Green was performed to ensure equal loading and transfer of protein. Then membranes were rinsed with Tris-buffered saline with Triton X-100 and blocked with 5% BSA for 1 h. Western blotting was performed with listed primary antibodies overnight at 4°C. Horseradish peroxidase-conjugated secondary antibodies (concentration 1:5000) were then added to the membrane in 5% milk. After washing for three times for 10 minutes each, Pierce ECL substrate was added to the membrane and developed using autoradiography film (Santa Cruz Biotechnology). Inhibitor experiments were performed in a similar fashion after overnight serum starvation. Starved cells were pre-treated with ruxolitinib (1 μM, Selleck Chemicals) or imatinib (1 μM, Selleck Chemicals) for one hour and then treated with either PDGF-BB (10 ng/mL, R&D Systems) or IFNγ (10 ng/mL, EMD Millipore) for 15 minutes prior to harvest.

### Phosphoproteomics and analysis

Fibroblasts were harvested and cultured as described above. At passage 1, cells were serum-starved overnight and then collected in 500 μL of cold PBS. Cells were then centrifuged at 1,000 rpm for 1.5 minutes at 4°C to form a cell pellet. Cells were shipped on dry ice to the IDeA National Resource for Quantitative Proteomics (Little Rock, Arkansas). Cells were purified by choloroform/methanol extraction and proteins were digested with trypsin and Lys-C to cleave at the C-terminal side of lysine and arginine residues. Then Tandem Mass Tag (11-plex) labeling was performed to individually identify each sample. TiO_2_ and Fe-NTA phosphopeptide enrichment was used to enrich for phosphopeptides. Then high-pH peptide fractionation was performed on enriched and un-enriched samples containing 18 fractions each. Fractions were injected into the Orbitrap Eclipse (TMT MS3 for 60 minutes gradient per fraction) for LC/MS analysis and identified by using MaxQuant. For analysis, all detected peptides, their respective log2 intensities, and the nominal p-values were used as inputs. To filter significance, a cut-off value of p < 0.05 was used for both Gene Set Enrichment Analysis and Ingenuity Pathway Analysis. The gene set m5.all.v2025.1.Mm.symbols.gmt was used as a reference. Permutations were limited to 1000 and compared the unnormalized expression of protein (converted to gene names) from P583R to WT. By converting protein to gene name, it allowed for use of the NCBI gene ID as the mouse collection Chip. RStudio version 2024.12.1+563 was used to generate visualization plots. Both proteomics and phosphoproteomics data were analyzed using libraries DEP, SummarizedExperiment, dplyr, tibble, readxl. Heatmaps were generated from the R package pheatmap.

### Statistical analysis

Data shown in figures are represented as mean ± standard deviation (weight curves) or mean ± SEM (all other data). Statistical significance between experimental groups was determined using GraphPad Prism 10 by unpaired Student’s t-test with standard deviation assumed to be equal between experimental groups. For more than two experimental groups, two-way ANOVA with multiple comparisons and Tukey correction was used for weight data analyses. Ordinary one-way ANOVA was performed with multiple comparisons and Tukey correction for bone analyses, skin thickness quantification, and qPCR data. The threshold for significance was considered as a p < 0.05. In proteomics and phosphoproteomics analyses, the threshold used for significance was log_2_ fold change >1 and a p value < 0.05.

## ACKNOWLEDGEMENTS

JHK was supported by NIH/NIDCR predoctoral fellowship F31-DE033289. This research was supported by R01-AR073828 and R01-AR080896 (LEO), R35-GM142786 (WLB), R03-DE034789 (HRK), by the Oklahoma Center for Adult Stem Cell Research (a program of TSET), and by grants from the Oklahoma City-based Presbyterian Health Foundation to LEO.

## AUTHOR CONTRIBUTIONS

JHK and LEO conceived and designed the study. JHK performed all experiments and analysis. HRK contributed to skeletal and bioinformatics analysis. WLB provided reagents and contributed to the creation of the mouse model. JHK wrote the manuscript. LEO supervised the research and edited the manuscript.

---

**Supplementalary Figure 1.**
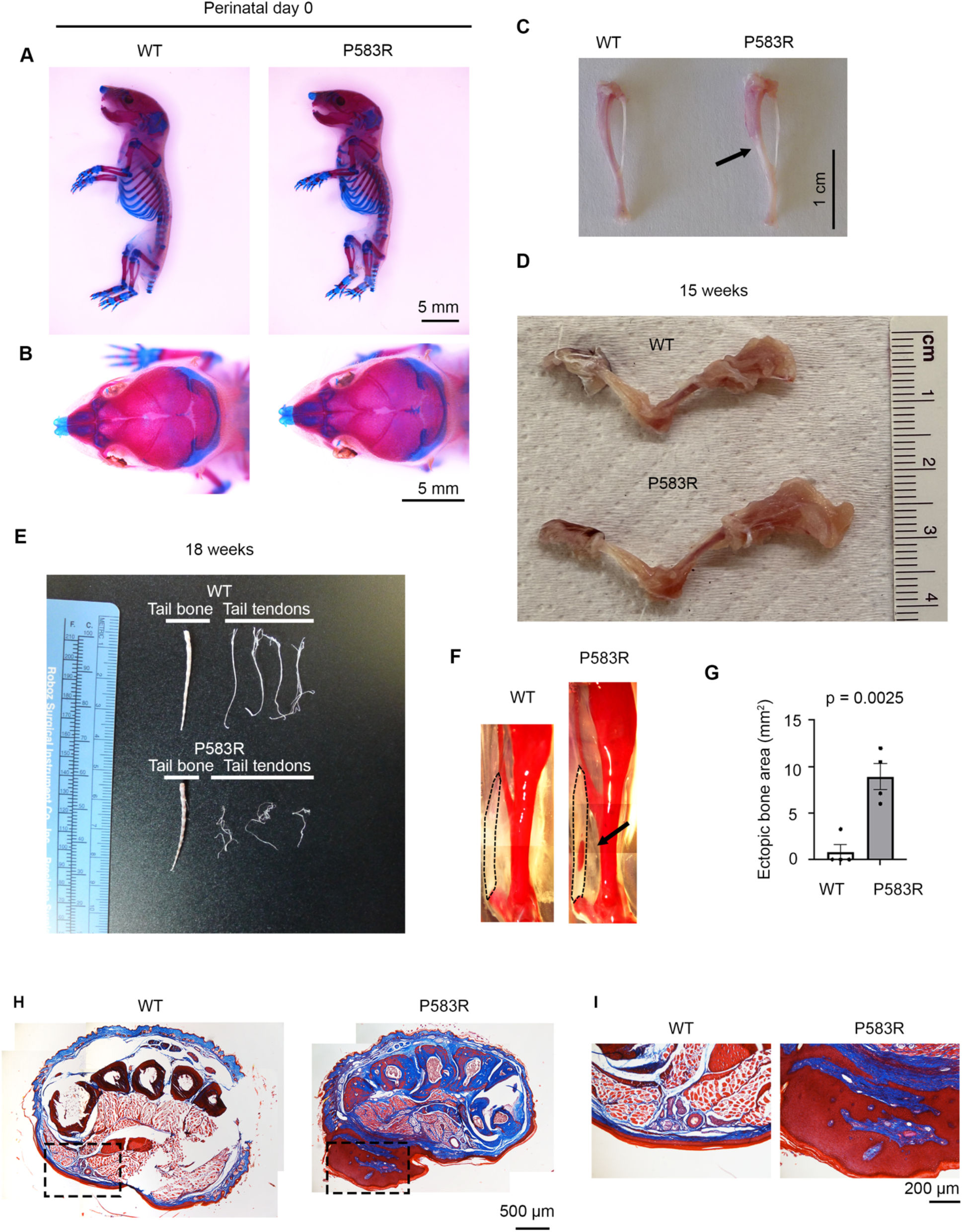
Skeletal phenotypes in P583R mice. (A-B) Whole mount Alizarin Red staining at postnatal day 0. (C) Dissected female tibias showing increased length of P583R at 15 weeks and absence of distal bone marrow (arrow). (D) Dissected forelimbs showing increased length of P583R at 15 weeks. (E) Isolated tail tendons at 18 weeks. (F) Alizarin Red staining of hindlimbs of WT and P583R mice at 9 weeks. Black arrow indicates heterotopic ossification in the Achille’s tendon, which is outlined with dotted line. (G) Quantification of ectopic bone area. Error bars indicate mean +/-SEM, unpaired Student’s t-test. n = 4 mice per genotype, each dot represents one mouse. (H) Masson’s Trichrome staining of paws from WT and P583R mice at 26 weeks. Dotted box area shown in (I). (I) Zoom in images of inferior portion of paw with a hyperkeratotic lesion in P583R.

**Supplementary Figure 2.**
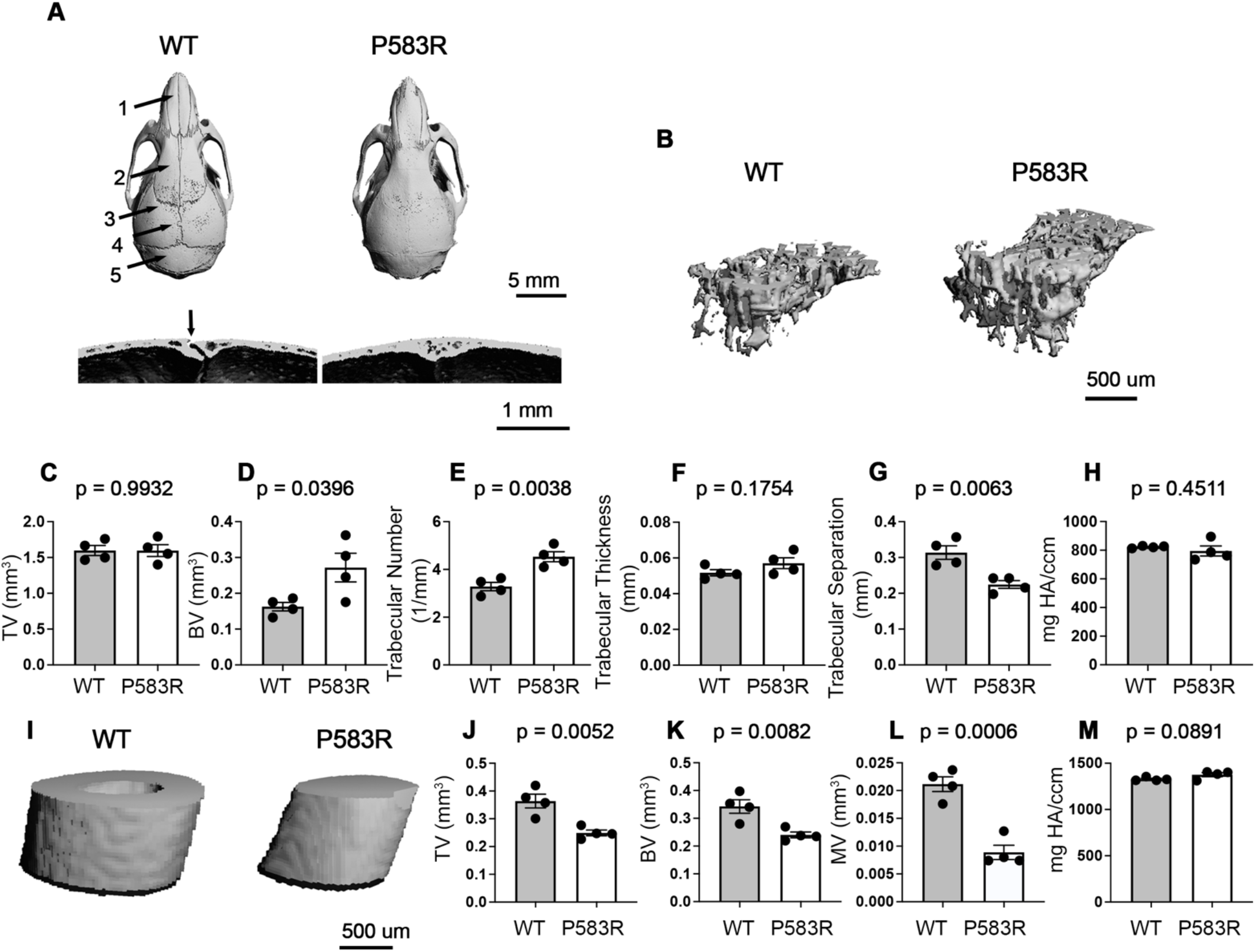
Skeletal Overgrowth in P583R females. (A) μCT images of female skulls at 15 weeks showing craniosynostosis of all P583R cranial sutures. Arrows indicate sutures. 1 = internasal suture, 2 = interfrontal suture, 3 = coronal suture, 4 = sagittal suture, 5 = lambdoidal suture. Arrow in the cross section view indicates patent lambdoidal suture in WT, which is closed in P583R. (B) 3D reconstruction from μCT scans of 15-week tibias showing trabecular bone. (C) Trabecular bone total volume. (D) Bone volume of trabecular bone. (E) Trabecular number. (F) Trabecular thickness. (G) Trabecular separation. (H) Trabecular bone mineral density. (I) 3D reconstruction from μCT scans of 15-week tibias at the point of fibula attachment. (J) Cortical bone total volume. (K) Bone volume of cortical bone. (L) Cortical bone medullary volume. (M) Cortical bone mineral density. Error bars indicate mean +/-SEM, unpaired Student’s t-test, n = 4 mice per genotype, each dot represents one mouse.

**Supplementary Figure 3.**
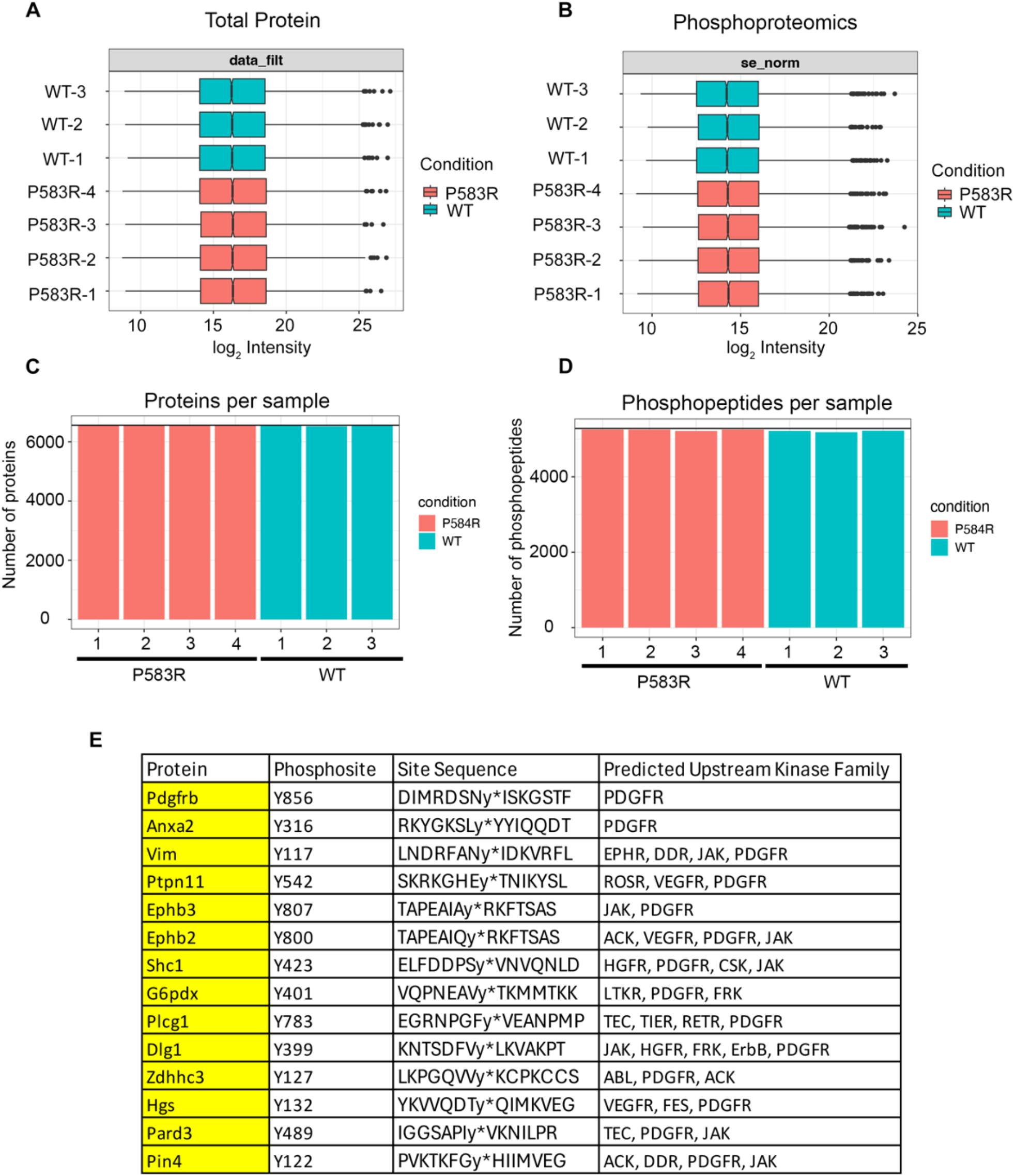
Phosphoproteomics identifies upregulated tyrosine phosphorylation on potential PDGFRβ substrates. (A) Normalized intensity values of total proteins from WT and P583R. (B) Normalized intensity values of phosphopeptides from WT and P583R. (C) Total number of detected proteins in P583R and WT samples. (D) Total number of detected phosphopeptides in P583R and WT samples. (E) List of differentially tyrosine-phosphorylated proteins predicted to have PDGFR as an upstream kinase. Predictions based on motif analysis from the kinase library^30^.

**Supplementary Figure 4.**
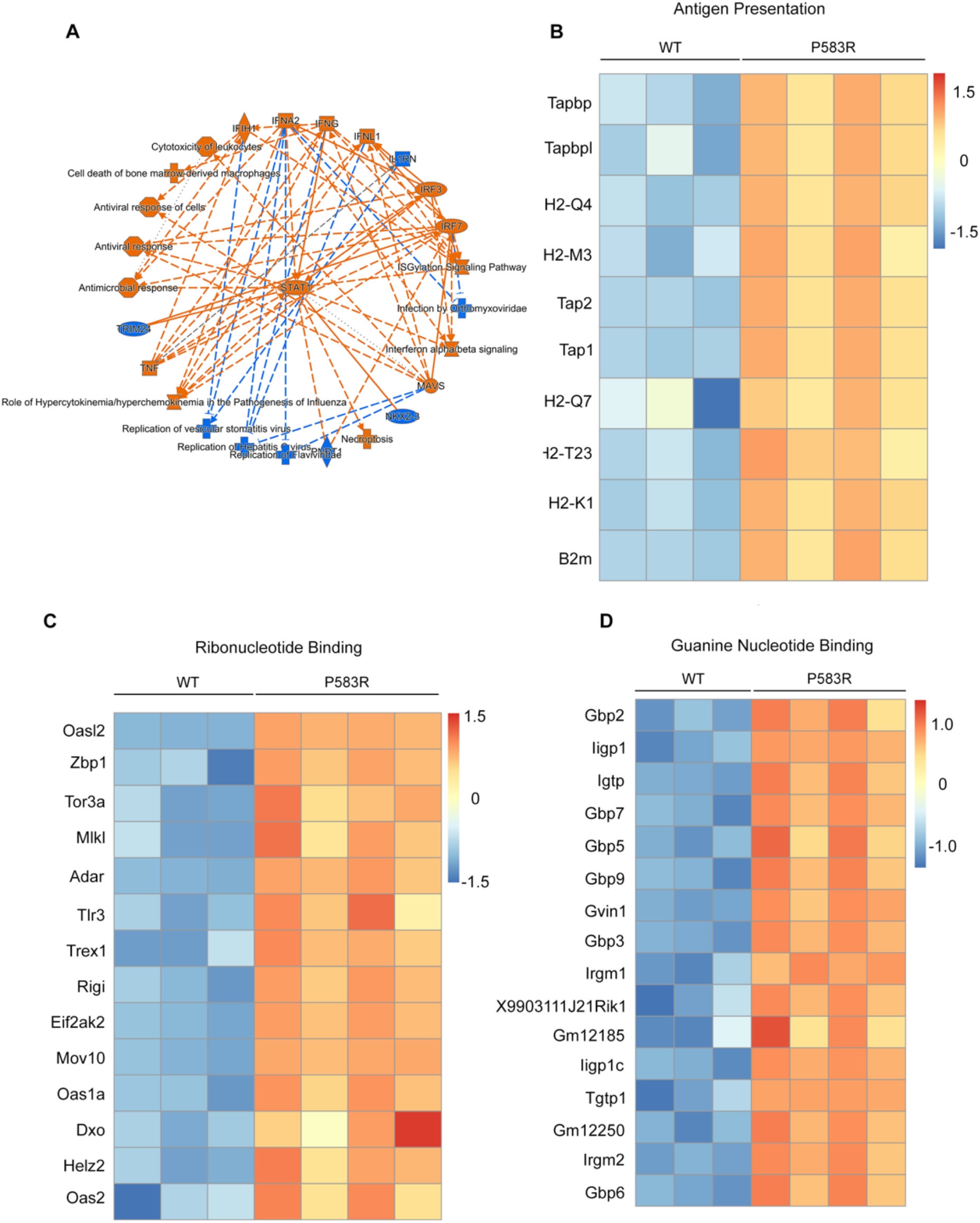
P583R induces hyperactivation of pro-inflmmatory signals, antigen presentation, and pathways related to nucleotide binding. (A) IPA graphical summary of pathways linked to STAT1 based on differentially expressed proteins. (B) Heatmap of protein expression comprising the “Antigen presentation” category obtained from GSEA. (C) Heatmap of protein expression comprising the “Ribonucleotdie binding” category obtained from GSEA. (D) Heatmap of protein expression comprising the “Guanine nucleotide binding” category obtained from GSEA. All cutoff values used for significance include log_2_ normalized intensity values > 1 and p < 0.05.

**Supplementary Figure 5.**
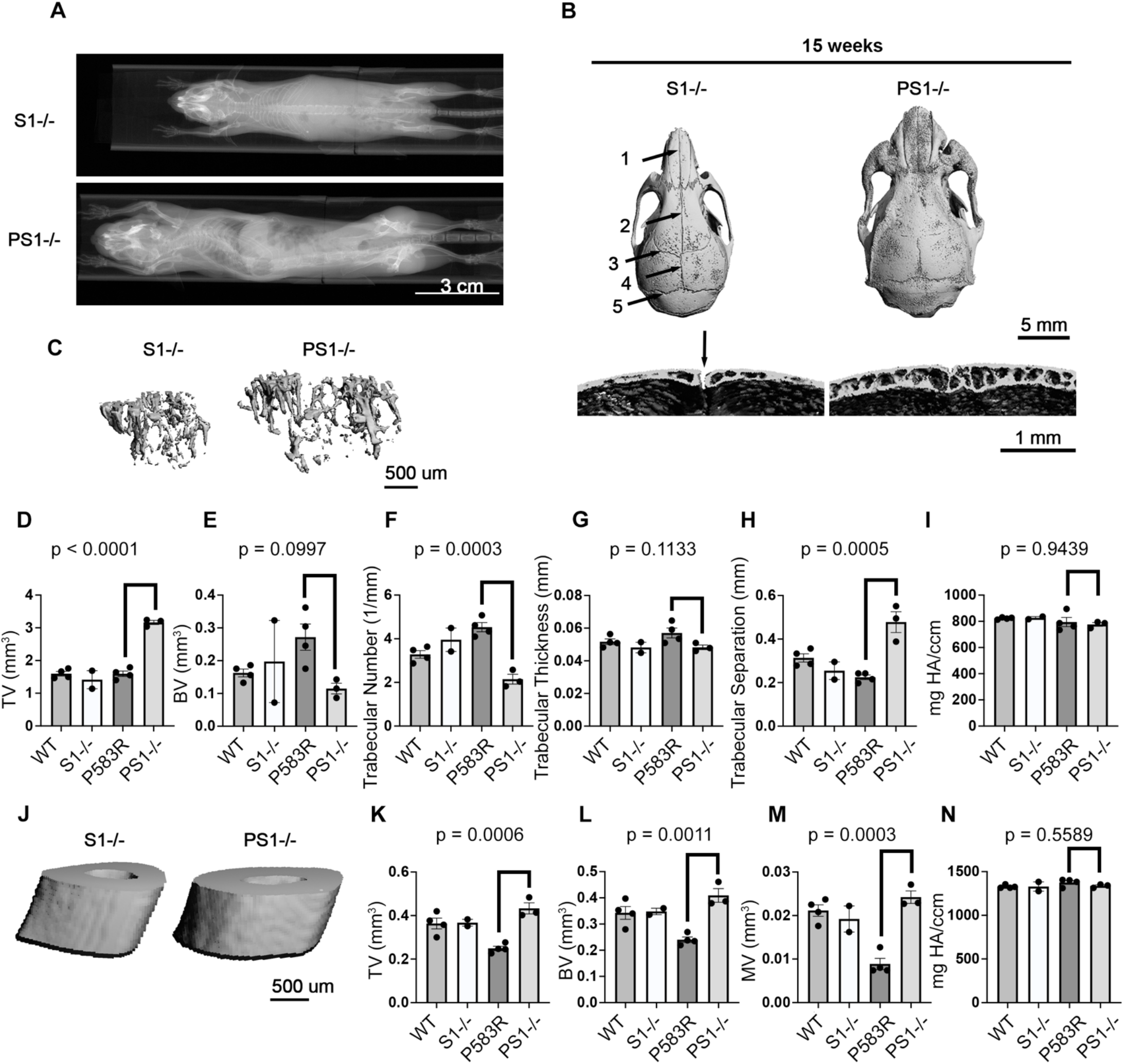
Exacerbated skeletal overgrowth in PS1^-/-^ females. (A) Whole body X-rays showing increased length and scoliosis of PS1^-/-^ females at 15 weeks. (B) μCT images of female skulls at 15 weeks. Arrows indicate sutures. 1 = internasal suture, 2 = interfrontal suture, 3 = coronal suture, 4 = sagittal suture, 5 = lambdoidal suture. Arrow in the cross-section view indicates patent lambdoidal suture in S1^-/-^, which is closed in PS1^-/-^. (C) 3D reconstruction from μCT scans of 15-week tibias showing trabecular bone. (D) Trabecular bone total volume. (E) Bone volume of trabecular bone. (F) Trabecular number. (G) Trabecular thickness. (H) Trabecular separation. (I) Trabecular bone mineral density. (J) 3D reconstruction from μCT scans of 15-week tibias at the point of fibula attachment. (K) Cortical bone total volume. (L) Bone volume of cortical bone. (M) Cortical bone medullary volume. (N) Cortical bone mineral density. (C-N) All quantifications from female mice at 15 weeks. Data for WT and P583R are the same as Supplementary Figure 2C-H, J-M. Error bars indicate mean +/-SEM, one-way ANOVA with multiple comparisons and Tukey’s correction, n = 2-4 mice per genotype, each dot represents one mouse.

